# In situ spatial reconstruction of distinct normal and pathological cell populations within the human adrenal gland

**DOI:** 10.1101/2023.06.12.544676

**Authors:** Rui Fu, Kathryn Walters, Katrina Koc, Amber Baldwin, Michael Clay, Katja Kiseljak-Vassiliades, Lauren Fishbein, Neelanjan Mukherjee

## Abstract

The human adrenal gland consists of concentrically organized functionally distinct regions responsible for hormone production. Dysregulation of adrenocortical cell differentiation alters the proportion and organization of the functional zones of the adrenal cortex leading to disease. Current models of adrenocortical cell differentiation are based on mouse studies, but there are known organizational and functional differences between human and mouse adrenal glands. This study aimed to investigate the centripetal differentiation model in the human adrenal cortex and characterize aldosterone-producing micronodules (APMs) to better understand adrenal diseases such as primary aldosteronism. We applied spatially resolved *in situ* transcriptomics to human adrenal tissue sections from two individuals and identified distinct cell populations and their positional relationships. The results supported the centripetal differentiation model in humans, with cells progressing from the outer capsule to the zona glomerulosa, zona fasciculata, and zona reticularis. Additionally, we characterized two APMs in a 72-year-old female. Comparison with earlier APM transcriptomes indicated a subset of core genes, but also heterogeneity between APMs. The findings contribute to our understanding of normal and pathological cellular differentiation in the human adrenal cortex.

## INTRODUCTION

The human adrenal gland consists of an outer cortex responsible for steroid hormone biosynthesis and an inner medulla responsible for catecholamine synthesis. The cortex is concentrically arranged into histologically and functionally distinct regions including the outer capsule, zona glomerulosa (zG), zona fasciculata (zF), and zona reticularis (zR) (1). The ability of cortical cells to self-renew and differentiate is crucial to normal adrenocortical homeostasis (2). Dysregulation of molecular pathways controlling cortical cell differentiation dynamics and/or hormonal secretion, leads to human diseases including primary aldosteronism and adrenocortical carcinoma, among others. The current centripetal differentiation model posits that adult stem and progenitor cells in the capsule/sub-capsular region differentiate into zG cells that further differentiate into zF cells and then zR cells (3–5). This model is based on experiments from mouse adrenal glands, which differ in at least two cell populations and functions from human adrenal glands (4). Nevertheless, these and other recent studies have found that the organization of layers and cell populations in the adrenal gland are more dynamic and heterogeneous than previously known.

An important example of cellular heterogeneity within histologic zones of the human adrenal cortex is the discovery of aldosterone-producing micronodules (APMs), formerly called aldosterone-producing cell clusters (6). APMs are defined as CYP11B2-positive cell clusters that are not discernible from surrounding cells of the capsule and zG by hematoxylin-eosin staining (7). APMs are associated with autonomous aldosterone production, and a subset of APMs may be precursors to aldosterone-producing adenomas (APA) (8). Although mouse models of APA exist, none of the models to date produce APMs, suggesting potential alternative pathways and highlighting the need to work with human tissue samples (9). Interestingly, APMs can be found in normal human adrenal tissue samples and they increase in frequency with age (10,11). Understanding the molecular etiology of APMs, as well as APAs, is critical in understanding primary aldosteronism (PA), an underdiagnosed treatable secondary cause of hypertension. In fact, there has been a recent conceptual shift toward recognizing primary aldosteronism as a continuum of autonomous aldosterone production that exists with varying severity even in normotensive individuals and all the way to severely hypertensive individuals (12,13). Unrecognized primary aldosteronism leads to cardiovascular disease, myocardial infarction, and stroke.

The overarching goal of this study was to identify supporting evidence for the centripetal differentiation model in the human adrenal cortex and identify pathways involved in APM development to better understand the etiology of primary aldosteronism. To advance our limited understanding of human adrenocortical cell differentiation and heterogeneity, we need to understand how cell populations within the human adrenal gland self-renew and differentiate. Preserving the spatial relationship between cells and using a global unbiased approach is a critical initial step. In this report, we apply spatially resolved *in situ* transcriptomics to human adrenal tissue sections to better understand the pathways controlling adrenocortical differentiation. To further investigate cellular heterogeneity, using APMs as an example, we characterized the transcriptomes of APMs compared with neighboring zG cells to identify markers able to discriminate between the two. Together, these data provide foundational knowledge to enhance our understanding of both normal and pathological cellular differentiation in the human adrenal cortex.

## RESULTS

### Determining the spatial relationships between human adrenal cell populations

Visium 10x spatial transcriptome analysis was performed on four different normal adrenal sections from two individual deceased donors (three sections from a 31-year-old female and one section from a 72-year-old female). Harmony was used to integrate the transcriptome data from the four sections (14) (**Supplemental Figure 1A, B**). On average, ∼3,000 genes per spot were detected for every section (**Supplemental Figure 1C**). The UMAP shows the twelve distinct and reproducible cell populations identified based on similarity in gene expression (**Figure 1A and Supplemental Figure 1D)**. Projecting the normal adrenal gland cell populations transcriptome back to the H&E stained tissue section (**Figure 1B, left**) revealed the expected concentric organization remarkably consistent with the histology (**Figure 1B, right**). For example, the expression of steroid hormone metabolism genes was highest in the cortical cells determined by visual mapping (**Figure 1C, top left**) and assigned cortical zones on UMAP (**Figure 1C, bottom left**). Likewise, the expression of amine derived hormone gene set was highest in the medullary cells (**Figure 1C, top and bottom right**). This observation was true for all four tissue sections (**Supplemental Figure 1E**). Next, we assessed the relative contribution of different cell populations to the human adrenal gland based on expression pattern. The adrenal cortex had 53% of the cellular contribution, whereas 17% was medulla, 10% was a mixture of cortex and medulla, and the remaining 21% was comprised of fibroadipose tissue, blood vessels or peripheral nerve cells (**Figure 1D**). As expected, genes involved in aldosterone, cortisol, and androgen production were enriched in zG (*CYP11B2*), zF (*CYP11B1*), and zR (*SULT2A1*), respectively (**Figure 1E)**. We identified known markers of the capsule such as *RSPO3*, as well as known markers of the zona glomerulosa such as *WNT4*. And, *PNMT* and *TH*, which are crucial for catecholamine production, were specifically expressed in the medulla. These results validate our cell population assignments.

**Figure 1.**
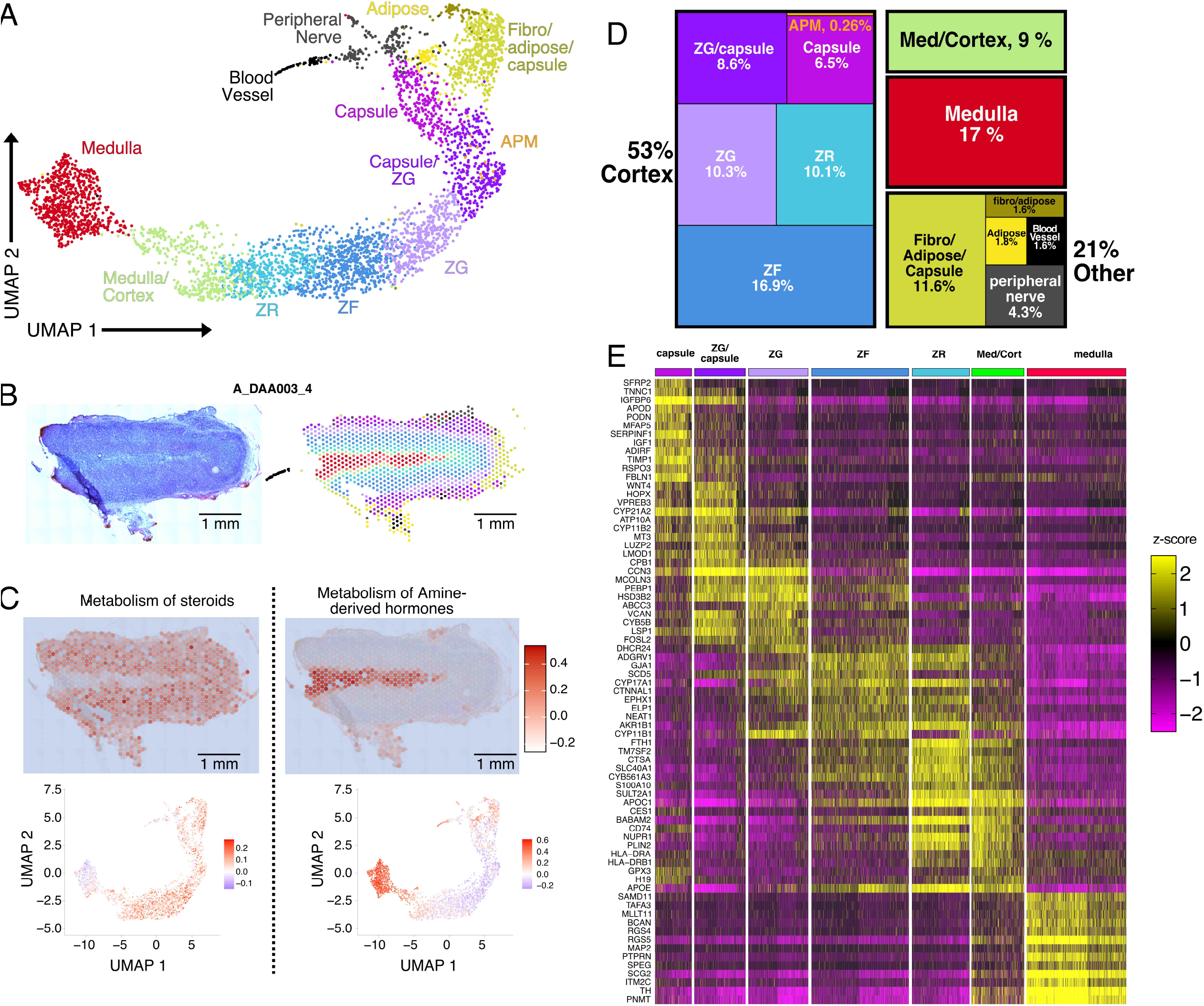
In situ reconstruction of distinct cell populations within the human adrenal gland. A) UMAP projection of spots from all sections following harmony integration color-coded by cell population assignment. B) H&E staining (left) of a representative donor adrenal section and in situ spots color-coded by cell population assignment (right). C) Expression of steroid metabolism (left) and amine-derived hormone (right) genes in situ (top) and on the UMAP projection (bottom). D) Treemap of the percentage of spots assigned to specific cell populations. E) Heatmap of the z-scores for the top differentially localized genes.

### The centripetal differentiation model in the human adrenal cortex

Next, we examined if the centripetal differentiation model based on mouse adrenal glands (3,5) was recapitulated in human adrenal glands. The developmental progression of the identified cortical cell populations was inferred using diffusion pseudotime analysis (DPT) (15). This analysis indicated that capsule to zG to zF to zR is the order of differentiation and transition between cell types of the adrenal cortex (**Figure 2A**). Classic zone-specific markers for zG (*CYP11B2*), zF (*CYP11B1*), and zR (*SULT2A1*) support the observed order of cell population transitions (**Figure 2B**). Furthermore, the peak enrichment of *RSPO3* in the pseudotime regions corresponding to the capsular cells precedes the peak enrichment of *WNT4* in the pseudotime regions corresponding to the zG cells (**Figure 2B**). Interestingly, both *RSPO3* and *WNT4* genes are members of the WNT signaling pathway and have been shown to regulate the balance between self-renewal and differentiation in the mouse adrenal gland capsule and zG (**Figure 2B**) (16,17). We also identified novel genes with restricted expression patterns associated with the capsule (*IGFBP6* and *FBNL1*) and capsule/zG (*LMOD1* and *SFRP2*), respectively (**Figure 2B**). The pseudotime values for individual genes were consistent with their expression in cortical cell populations (**Supplemental Figure 2A**). The observed transition between cell populations and the spatially restricted expression of key adrenocortical regulators of self-renewal and differentiation suggest that the centripetal differentiation model determined in mice is consistent in normal human adrenals.

**Figure 2.**
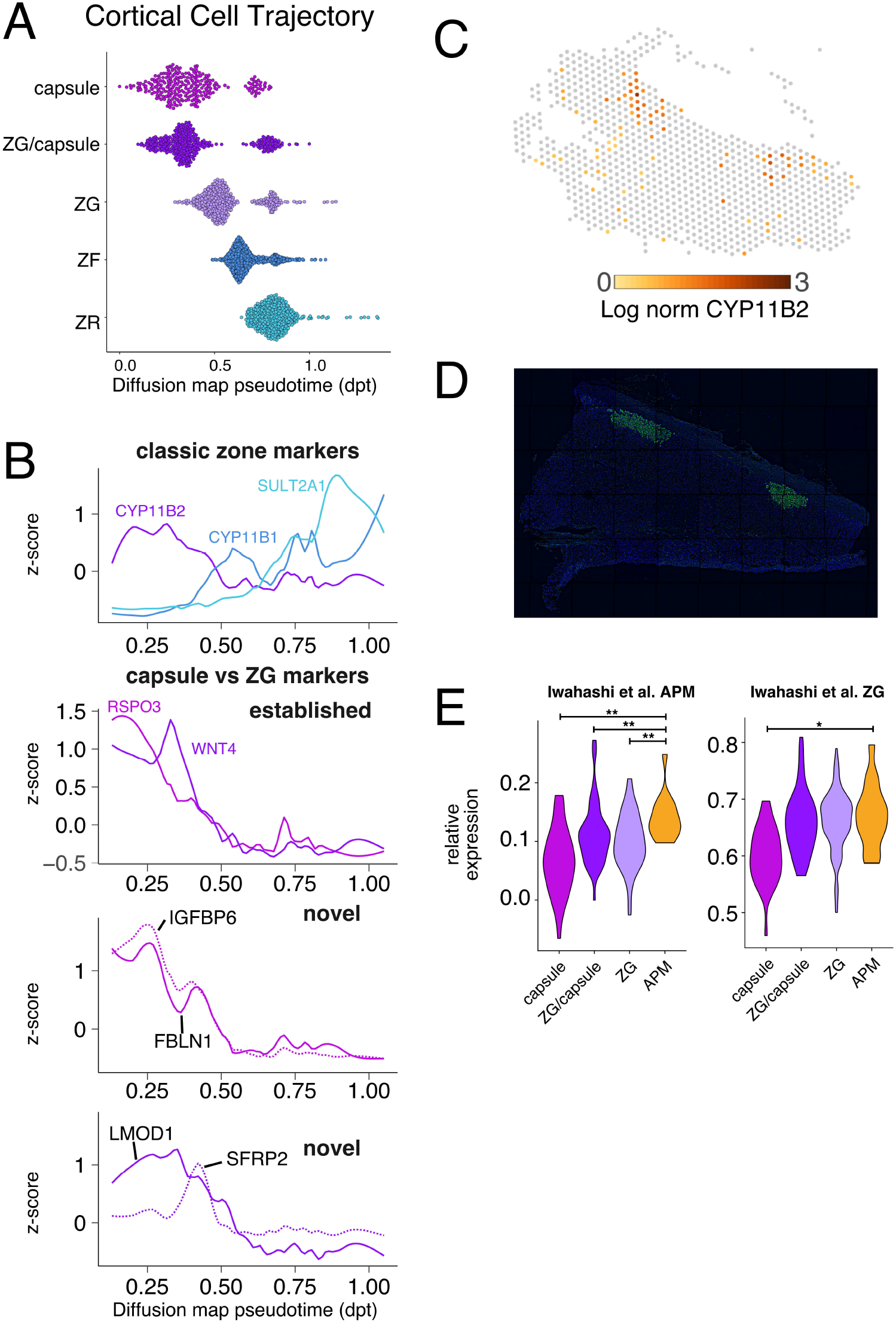
In situ reconstruction of distinct cell populations within the human adrenal gland. A) Dotplot of the distribution of diffusion map pseudotime (dpt) values for each adrenocortical cell population. B) Line plot of expression z-scores versus dpt values for genes colored by their respective cell population. C) *CYP11B2* mRNA expression for adrenal section corresponding to 72-year-old female. D) *CYP11B2* protein staining (green) and DAPI (blue) for adjacent adrenal section corresponding to 72-year-old female. E) Violin plot of relative expression levels for genes associated with either APM (left) or zG (right) from Iwahashi et. al (*p <.05, **p <.01 Wilcoxon test).

### The transcriptome of aldosterone-producing micronodules (APMs)

Aldosterone-producing micronodules (APMs) are an important example of spatially restricted cell heterogeneity in the adrenal cortex. APMs are defined by clustered high protein expression of the CYP11B2 enzyme (aldosterone synthase). In our samples, the *CYP11B2* mRNA (not shown) and protein expression (**Supplemental Figure 2B**) in the 31-year-old donor exhibited a typical continuous pattern across most cells of the zG layer. However, *CYP11B2* mRNA expression was discontinuous and localized to regional densities in the adrenal gland section from the 72-year-old donor (**Figure 2C)**. CYP11B2 protein by immunofluorescence on an adjacent section similarly revealed two positive staining regions consistent with multiple APMs (**Figure 2D**). Interestingly, the spots containing APMs were most similar in gene expression to capsule/zG cell populations (**Figure 1A UMAP**). Expression signatures of APMs have been suggested in previous studies either through analysis by single nuclei RNA-seq or laser capture microdissection. The APM gene signature identified using single-nuclei RNA-seq (18) was significantly enriched in our APM cell populations relative to other cortical cell populations (**Figure 2E, right**). We did not observe a similar significant enrichment using gene sets identified using laser capture followed by microarray (19) (**Supplemental Figure 2C**). The zG gene signature from Iwahashi et. al. had similar expression between our APM and zG populations, further supporting that the specificity of the APM signature similarity. We observed that *STAR* expression was higher in our APM cells compared with other cortex cells, indicating they have ample machinery of the rate-limiting step enzyme to produce excess aldosterone (**Supplemental Figure 2C**), consistent with the clinical picture in primary aldosteronism. Interestingly, each of the steroidogenic zone cell populations had a subpopulation with higher *STAR* expression, which may reflect further heterogeneity with respect to zone-specific hormone production. Altogether, we characterized the transcriptome of APMs in a 72-year-old human adrenal tissue section, which appears to have higher steroidogenic potential (i.e. high *STAR* expression) and is enriched for genes identified in prior studies of APM-expression signatures.

## DISCUSSION

Mouse models are powerful tools to understand human development and disease. However, species-specific differences and the lack of human disease models are major barriers, especially for adrenal cortical cell differentiation, which still requires the development of bona fide human stem cell differentiation models (3,5). Thus, studying human tissue is an essential resource to understand adrenal cell differentiation and the dysregulation of that process leading to disease. Here we applied spatially resolved transcriptomics to normal adrenal tissue sections from a 31-year-old female and a 72-year-old female. In spite of the fact that the mouse adrenal cortex does not have a well-defined zona reticularis (20), our data clearly suggest that the basic mechanism of the centripetal differentiation model is conserved in humans and mice, shown by the pseudotime analysis. Furthermore, we found that WNT pathway members required for proper cell differentiation, *RSPO3* and *WNT4*, have precisely the same spatially restricted expression in capsule and zG cells, respectively (17,21). Another interesting spatially restricted marker is *SRFP2*, Secreted Frizzled-Related Protein, which is a soluble modulator of WNT signaling expressed in the same cell populations as *WNT4*. Interestingly, SRFP2 is downregulated in APA and loss of SRFP2 promotes aldosterone production through inhibition of WNT signaling in mice (22). We identify numerous markers of specific cell populations that will be a resource for the community.

We identified multiple APMs in the zG region of the adrenal from a 72-year-old female. When comparing two previously defined APM gene signatures, only one of them was enriched in our APM cell population (18,19). The lack of concordance between all signatures could be due to technical limitations or indicate substantial heterogeneity exists between APMs from different individuals. Increasing the sample size, including sections from males, and continuing to span wide age ranges will be crucial to better understand APM heterogeneity and determine how it relates to PA and disease progression. The spot size of the Visium platform used in this study is 55 μm in diameter, which, therefore, includes contributions from multiple cells. We did not see obvious signatures for immune cell populations, likely because they are typically distributed throughout the tissue rather than in distinct spatial patterns of cortical cell populations. Moving forward, higher resolution and/or the use of deconvolution methods such as STdeconvolve (23) will enable a better assessment of the spatial positioning of immune cells. In summary, a deeper understanding of normal adrenal function will help identify changes underlying both the APA and neoplastic processes. Recent multi-omic technological developments combining spatially resolved DNA mutations (24) with RNA expression will be powerful for classifying and understanding adrenal pathologies.

## MATERIALS AND METHODS

### Adrenal gland processing

With IRB approval (COMIRB 15-0516), we worked with the Donor Alliance to obtain normal adrenal tissue attached to donor kidneys which would otherwise be discarded. Donors have already agreed to be organ donors and agreed to share donor tissue for research purposes. Normal adrenal glands were dissected and placed into histidine tryptophan ketoglutarate (HTK) solution. Within 2 hours of receiving the tissue, the gland was cut into sections for fresh frozen tissue and to create OCT blocks for storage. 10-micron sections of OCT blocks were cut and used for H&E to ensure a well-preserved tissue to use in further experiments. Adjacent sections were cut and used for spatial transcriptomics and immunohistochemical analysis.

### Spatial transcriptomics sample processing

Frozen samples were OCT embedded and sectioned at 10μm on a Cryostar NX70 cryostat (Thermo Fisher Scientific). Capture sections were fixed with methanol, stained with H&E, and imaged on an Evos M7000 (ThermoFisher) with brightfield settings. Capture sections were then permeabilized and processed to generate RNA libraries following the 10x Visium protocol. Libraries were sequenced to a depth of 60,000 read pairs per spot calculated from the image, on a Novaseq6000 (Illumina) sequencer with 151x151 bp runs.

### Spatial transcriptomics data analysis

Sequencing data were processed with Space Ranger (10x genomics, v1.2.1), followed by further analysis using the Seurat (v4.0.1) tool suite in R. Spots were lightly filtered to ensure the number of genes detected fall between 50 and 15000, and less than 50% of UMIs mapped to mitochondrial genes. After initial SCTransform normalization on each sample and principal component analysis on merged data, integration was performed with Harmony (v1.0) using 30 principal components and theta=2. UMAP dimension reduction and shared nearest neighbor clustering were carried out on 30 principal components, and clustering results at different resolution settings were explored through Clustree (v0.4.3) and 10x Cloupe browser visualizations. Specific cells overlaying H&E regions of interest were manually traced in 10x Cloupe browser and then exported to retrieve barcodes.

Differential ST cluster gene expression was defined by Wilcoxon test as implemented in Presto (https://github.com/immunogenomics/presto), with thresholds of adjusted p-value <= 0.001 and log2 fold change >= 0.5. Cell type identity of clusters was defined in 3 ways: (i) manual inspection of key markers, (ii) expert histological annotation by a pathologist of H&E stained slides, and (iii) Jaccard index calculation of ST marker gene overlap with previously reported scRNAseq markers from Huang et al. 2021 using Clustifyr. For medulla and zR/medulla we initially identified two cell populations, which we collapsed to single cluster based on a minimal number of differentially-expressed genes between the original two clusters. Per-cell gene set expression scoring was calculated through a R/rust reimplementation that speeds up the Seurat::AddModuleScore algorithm (https://github.com/raysinensis/SCoreRust). Pathways were defined by C2: curated gene sets in the Human MSigDB Collections release 7.5.1 (25). Pseudotime trajectory was inferred with the R package Destiny, designating zR cells as the tip of the diffusion branches. Z-scores for gene expression along pseudotime was calculated as 50 roughly equal cell number bins.

### Immunohistochemical analysis of tissue samples

Adrenal sections were fixed in 10% Neutral Buffered Formalin for 10 minutes. They were then blocked and permeabilized with CAS-Block + 0.2 TritonX (CAS-T) for 30min. Samples were stained overnight at 4C with anti-mouse CYP11B2 (26) diluted 1:1000 in CAS-T. The primary solution was rinsed with TBS+ 0.1% Tween20 (TBS-T) twice. Anti-mouse IgG Alexa Fluor® 488 was diluted 1:1000 in TBS-T and the sample was incubated at room temperature for 1 hour. Samples were rinsed with TBS-T and then mounted using VectaShield Vibrance™ Antifade Mounting Medium with DAPI. Slides were imaged with a 10x lens on a DeltaVision Elite Deconvolution Microscope and stitched together with DeltaVision software.

## Data availability

ST data have been deposited in the NCBI Gene Expression Omnibus (GEO) database and are publicly accessible through GEO accession number. A browsable internet resource of the adrenal ST data, including H&E histology, clusters and gene expression for all samples, is available at https://raysinensis.shinyapps.io/spatialshiny_adr/.

---

**Supplemental Figure 1.**
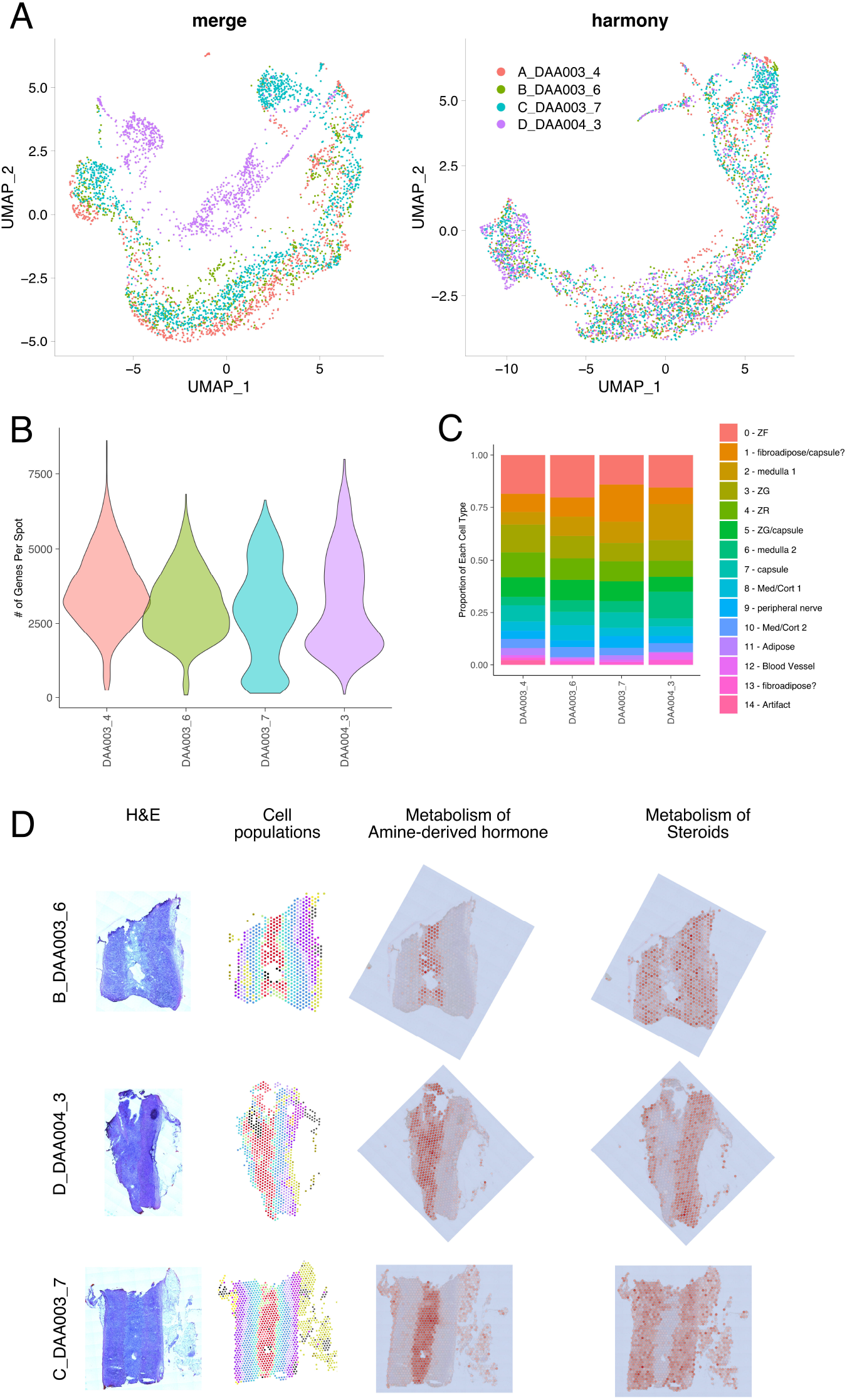
Processing, quality control, and integration of multiple tissue sections. A) UMAP projection of spots from each slide (right) and all sections following harmony integration (left). B) Violin plot of number of genes per section. C) Fraction of original cell population assignments per slide before collapsing similar cell populations. D) In situ visualization of H&E, cell populations, amine-derived hormone and steroid hormone genes, from left to right, respectively.

**Supplemental Figure 2.**
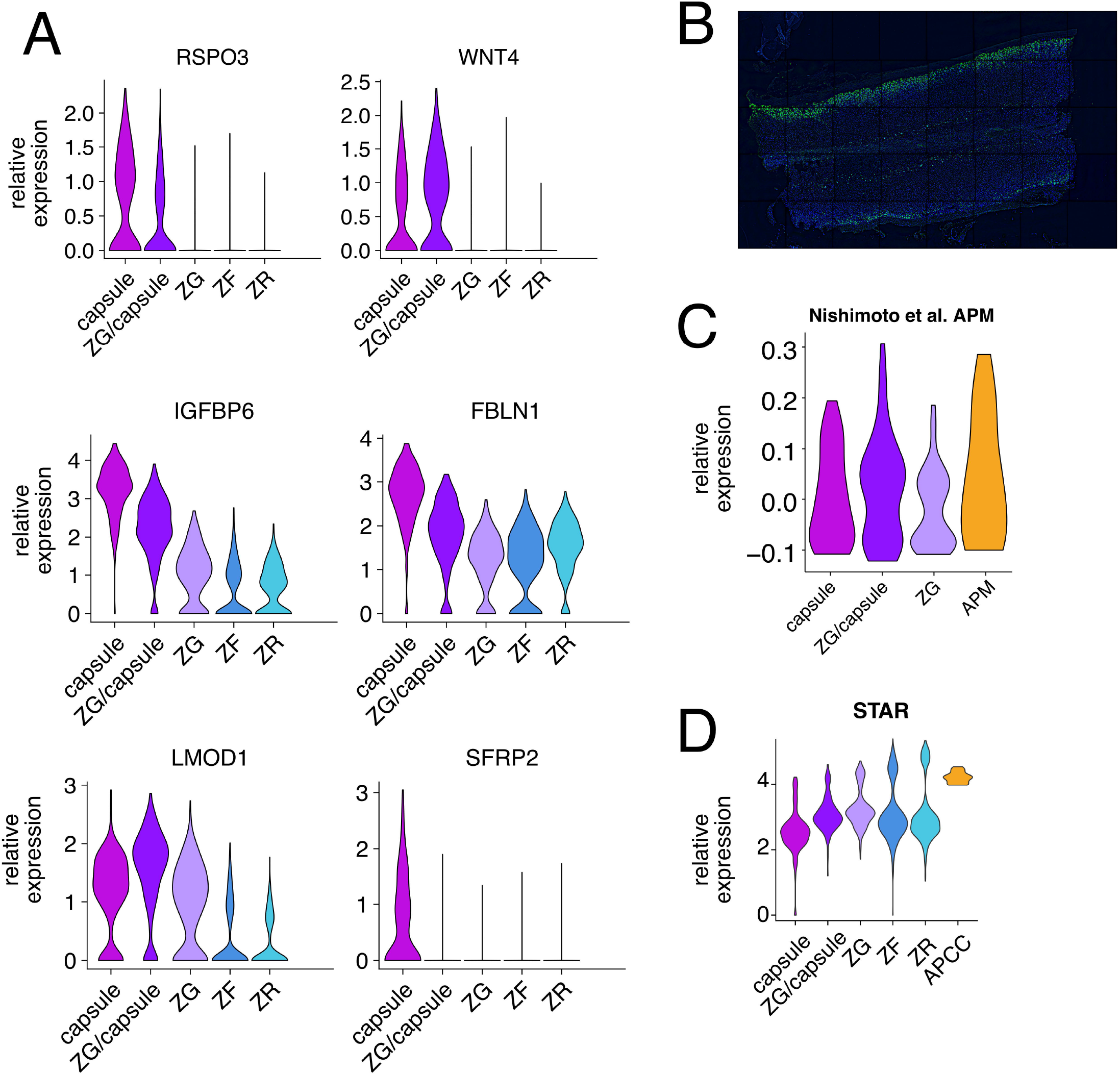
Characterizing mRNA and protein expression in adrenal section. A) Violin plot of relative expression levels for each cortical cell population for different genes. B) *CYP11B2* protein staining (green) and DAPI (blue) for adjacent adrenal section corresponding to 31-year-old female. C) Violin plot of relative expression levels for genes associated with APM from Nishimoto et. al. D) Violin plot of relative expression levels for each cortical cell population for *STAR*.

